# Genomic variations in paired normal controls for lung adenocarcinomas

**DOI:** 10.1101/153825

**Authors:** Li-Wei Qu, Bo Zhou, Gui-Zhen Wang, Ying Chen, Guang-Biao Zhou

**Affiliations:** Division of Molecular Carcinogenesis and Targeted Therapy for Cancer, State Key Laboratory of Membrane Biology, Institute of Zoology, Chinese Academy of Sciences, Beijing 100101; University of the Chinese Academy of Sciences, Beijing 100049

## Abstract

Somatic genomic mutations in lung adenocarcinomas (LUADs) have been extensively dissected, but whether the counterpart normal lung tissues that are exposed to ambient air or tobacco smoke as the tumor tissues do, harbor genomic variations, remains unclear. Here, the genome of normal lung tissues and paired tumors of 11 patients with LUAD were sequenced, the genome sequences of counterpart normal controls (CNCs) and tumor tissues of 513 patients were downloaded from TCGA database and analyzed. In the initial screening, genomic alterations were identified in the “normal” lung tissues and verified by Sanger capillary sequencing. In CNCs of TCGA datasets, a mean of 0.2721 exonic variations/Mb and 5.2885 altered genes per sample were uncovered. The C:G→T:A transitions, a signature of tobacco carcinogen N-methyl-N-nitro-N-nitrosoguanidine, were the predominant nucleotide changes in CNCs. 16 genes had a variant rate of more than 2%, and CNC variations in *MUC5B*, *ZXDB*, *PLIN4*, *CCDC144NL*, *CNTNAP3B*, and *CCDC180* were associated with poor prognosis whereas alterations in *CHD3* and *KRTAP5-5* were associated with favorable clinical outcome of the patients. This study identified the genomic alterations in CNC samples of LUADs, and further highlighted the DNA damage effect of tobacco on lung epithelial cells.

## Introduction

Comprehensive sequencing efforts over the past decade have confirmed that cancer is a disease of the genome^1^. Genomic alterations in cancers include point mutations, insertions and deletions (indels), structural variations, copy number alterations, epigenetic changes, and microbial infections. Abnormalities in oncogenes and tumor suppressors act as driver mutations to initiate the onset and progression of cancers. Technically, these alterations are identified by comparing the cancer genome sequence with a reference human genome and excluding those also found in normal controls (counterpart normal tissues or peripheral blood) and single nucleotide polymorphisms (SNPs)^2^. This strategy is effective to uncover somatic genomic alterations in tumor tissues. However, whether there is any genomic alteration in counterpart normal controls (CNCs) but not cancer tissues, remains unclear.

Lung cancer is the most common cause of cancer related mortality worldwide^3^ and can be divided into small cell lung cancer (SCLC) and non-small cell lung cancer (NSCLC). NSCLC comprises of lung adenocarcinoma (LUAD), lung squamous cell carcinoma (LUSC), and large cell carcinoma^4^. It is estimated that 90% of lung cancer deaths are caused by cigarette smoke^5^, whereas SCLC and LUSC are most strongly linked with smoking^4^. Since the late 1980s, LUAD has become the most common type (40%) of lung cancer that is related to cigarette smoke due to the changes in cigarette composition and filtering which favor adenocarcinoma histology^6^. There is also a proportion of LUADs that is not associated with smoking^4^, while air pollution represents another cause of lung cancer which induces somatic mutations in the genomes^7^. The cellular injury produced by smoking involves the whole respiratory tract^8, 9^; in the initial phase of lung carcinogenesis, injury is repaired by stem/progenitor cells, forming a clonal group of self-renewing daughter cells. Accumulation of genetic alterations in the premalignant field results in proliferation of the cells and expansion of the field, gradually displacing the normal epithelium and developing into a malignant neoplasm^10, 11, 12^. Genomic alterations of LUADs had been characterized and druggable targets had been unveiled^13, 14^. However, very few studies systematically dissect genomic mutations specifically occurred in “normal” lung tissues and their roles in carcinogenesis, and no study reports how the adjacent tissues escape from developing into malignant neoplasms.

In our work analyzing somatic genomic mutations in air pollution-related lung cancers^7^, we noted that when compared with the reference human genome, some variations were seen in counterpart normal lung tissues but not cancer samples (Table 1). To expand these observations, we further analyzed the Cancer Genome Atlas (TCGA) lung cancer genome data of 513 CNCs and counterpart tumor samples (Figure S1). Our results showed that the CNCs did harbor some genomic variations.

**Table 1.**
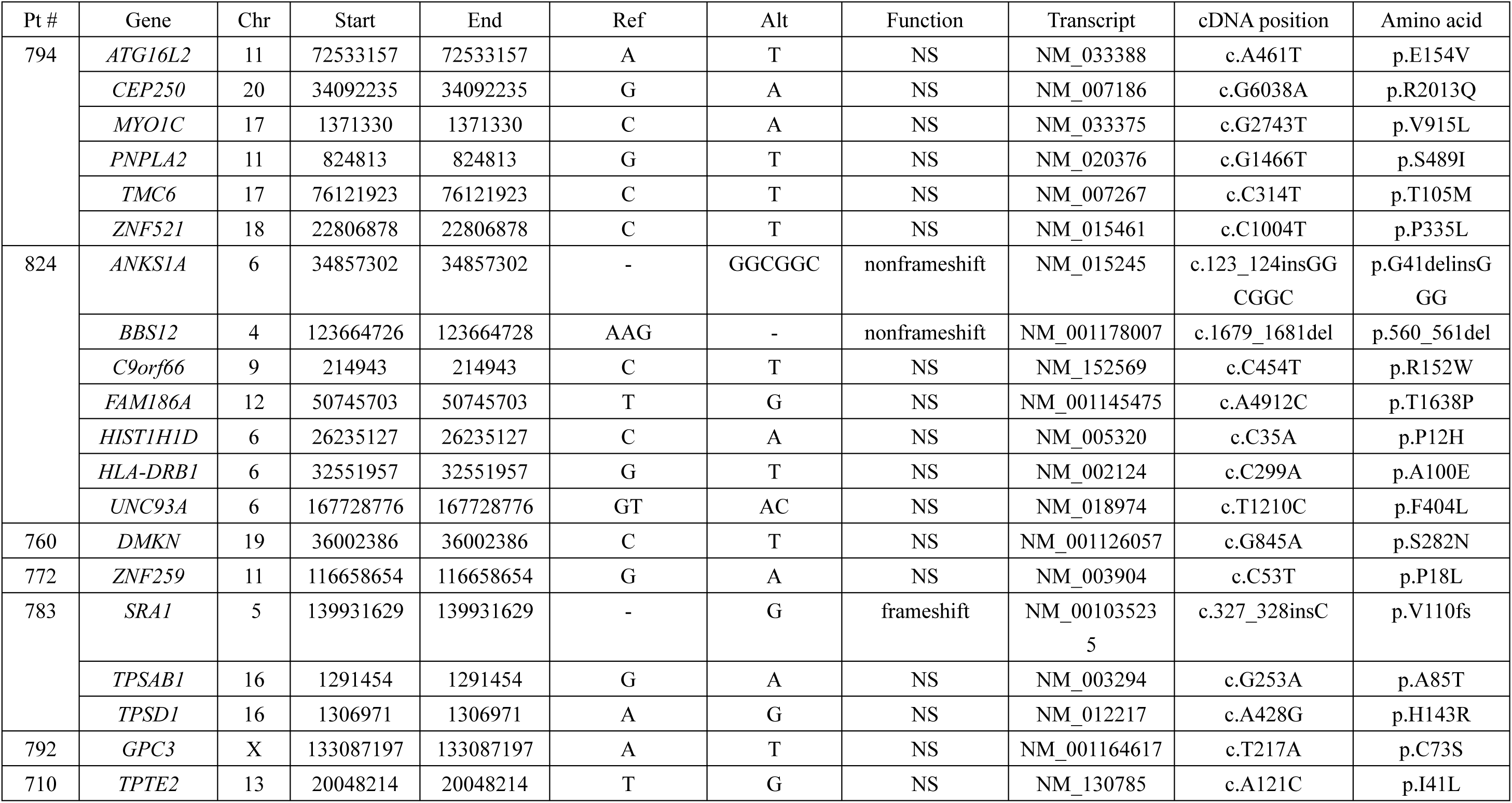
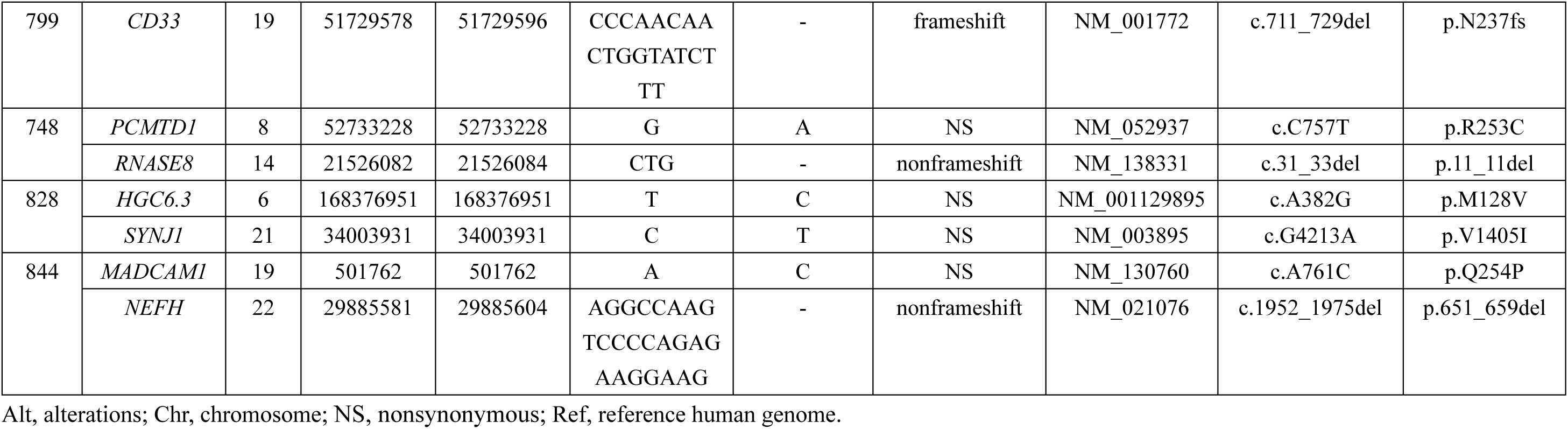
Genomic variations in CNCs of 11 patients with LUAD.

## Results

### Identification of CNC somatic genomic variations

The genomic DNAs of paired normal lung tissues and cancer tissues were sequenced to an average of 42.89× (30.07×–78.70×) coverage and 66.04× (range, 61.02×–74.64×) coverage, respectively. Twenty nucleotide substitutions and 7 small insertions and deletions (indels) were found in the 11 LUADs genomes (Tables 1). Variations in 6 representative genes in the normal lung tissues were validated by RT-PCR assays and subsequent sequencing (see Table S1 for sequence of primers), and the results confirmed the existence of CNC alterations (Figure 1). For example, the nucleotide at chr18: 22806878 of *ZNF521* of hg19 is C; but two peaks (C and T) of approximately equal peak height were seen in sequence of normal lung tissues, whereas a high peak of C and a low peak of T were detected in tumor samples of the patient (Figure 1A, left panel). This change might lead to P335L substitution in the encoded protein. Sequencing results using another set of primers confirmed the existence of T in normal lung rather than counterpart tumor sample of the patient (Figure 1A, right panel). Nucleotide T at this position of *ZNF521* was not found in dbSNP138 common germline variants. The nucleotide at chr17: 76121923 of *TMC6* of hg19 is C; but two peaks (C and T) of approximately equal peak height were seen in sequence of normal lung tissues, whereas a high peak of C and a low peak of T were detected in tumor samples of the patient (Figure 1B, left panel). This change might lead to T105M substitution in the encoded protein. Sequencing results using another set of primers confirmed the existence of T in normal lung rather than counterpart tumor sample of the patient (Figure 1B, right panel). Similarly, the normal lung tissues had a single nucleotide change in *CEP250* (Figure 1C), *C9orf66* (Figure 1D), and *HIST1H1D* (Figure 1E), compared to those in the tumor samples, hg19, and dbSNP138. Substitutions of GT by AC were seen in *UNC93A* in normal lung tissue of a patient with untreated LUAD (Figure 1F).

**Figure 1.**
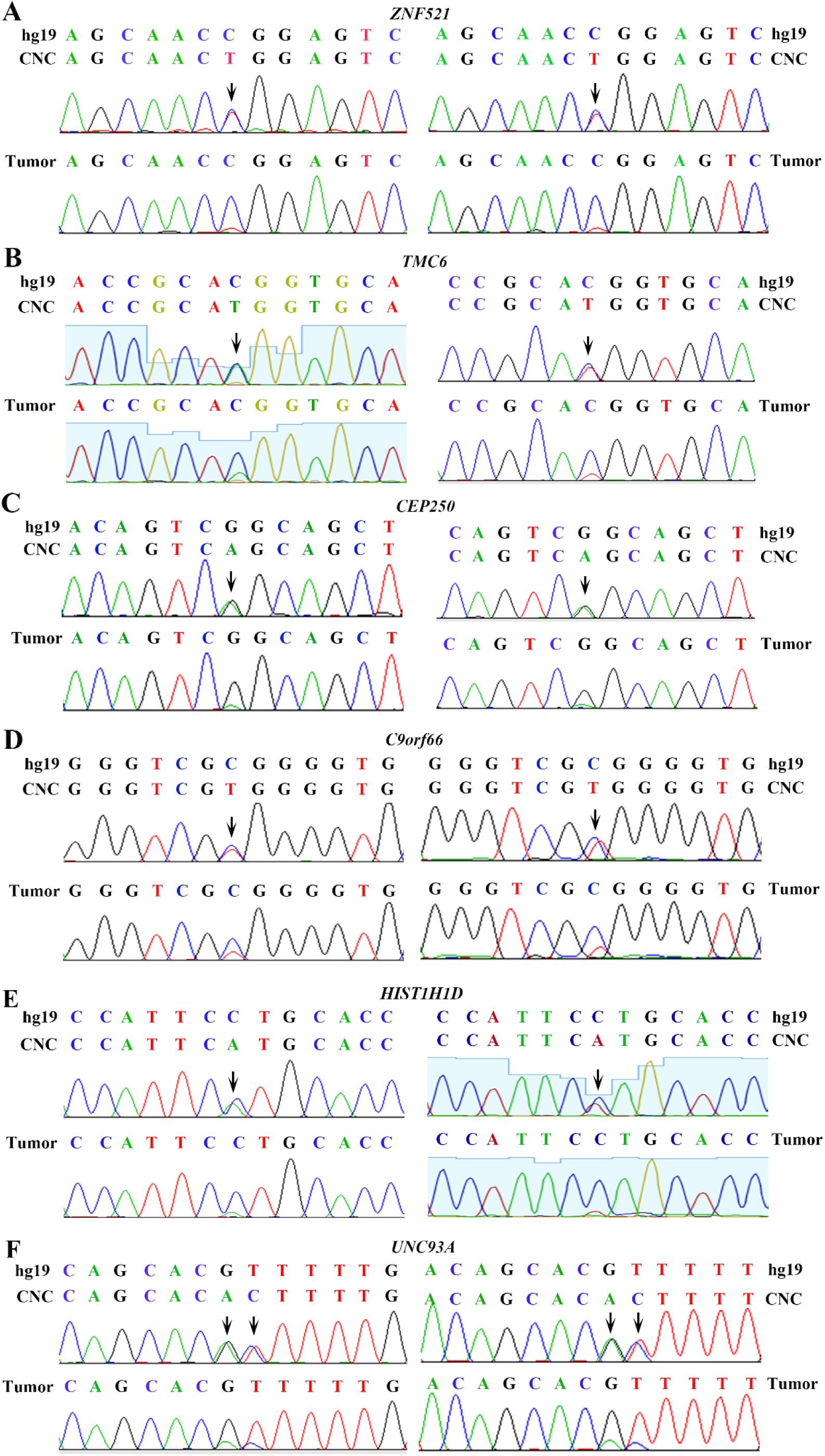
Validation of genomic variations in “normal” lung tissues. Polymerase chain reaction (PCR) and Sanger capillary sequencing were performed using genomic DNAs of the patients and primers listed in Table S1. (A) *ZNF521* in “normal” lung and tumor samples of the patient. Two sets of primers were used. (B) *TMC6* in “normal” lung and tumor samples of the patient. Two sets of primers were used. (C) *CEP250* in “normal” lung and tumor samples of the patient. Two sets of primers were used. (D) *C9orf66* in “normal” lung and tumor samples of the patient. (E) *HIST1H1D* in “normal” lung and tumor samples of the patient. (F) *UNC93A* in “normal” lung and tumor samples of the patient.

### Analyses of TCGA datasets

To expand the above observations, the genome sequence of cancer-control paired samples from 513 patients were downloaded from TCGA datasets and carefully analyzed. Among these patients (Table S2), 239 (46.6%) were males and 274 (53.4%) were females, and the median age was 66 years old (range, 33 – 88 years). Smoking history was available in 499 patients, among them 425 (85.2%) were current or reformed smokers (persons who were not smoking at the time of interview but had smoked at least 100 cigarettes in their life) and 74 (14.8%) were nonsmokers (not smoking at the time of interview and had smoked less than 100 cigarettes in their life). Adjacent normal lung tissues and peripheral blood were used as normal controls for 135 (26.3%) and 378 (73.7%) of the 513 lung adenocarcinomas, respectively.

Genomic variations were found throughout the counterpart control genomes. A mean of 0.2721 exonic variations/megabase (Mb) and 5.2885 variants/sample was recorded in the CNCs (Table 2). The normal lung tissues had approximately equal mutations to the blood cells (Table S3), while males and females had equal CNC mutations (Table S4). We compared the CNC mutations in nonsmokers, current smokers and reformed smokers, and found that smokers harbored more variations than nonsmokers, as reflected by variations/Mb (Figure S2A), altered genes/sample (Figure S2B), synonymous/nonsynonimous variations (Figure S2C), and indels/sample (Figure S2D). We also analyzed genomic mutations in tumor samples by comparing the cancer genome sequencing with reference and excluding those found in CNCs with the same calling criteria, and reported that tumor samples had much more mutations than CNCs (Table 2 and Figure S2, E-H).

**Table 2.**
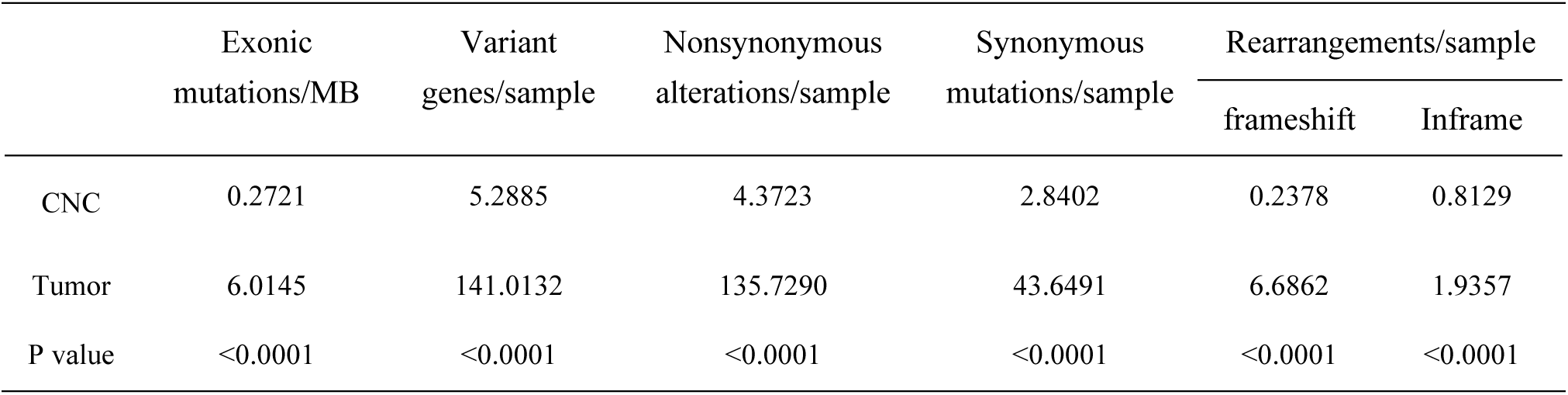
Genomic variations in CNCs and tumor samples of TCGA LUADs.

### Nucleotide substitutions in TCGA CNC samples

Tobacco smoke contains more than 20 lung carcinogens, e.g., nicotine-derived nitrosaminoketone (NNK)^5^ and polycyclic aromatic hydrocarbons (PAHs) which are associated with the C:G→A:T transversions in the genomes^7, 15, 16^. We analyzed the nucleotide substitutions in the genome, and found that the C:G→T:A and the A:T→G:C transitions were the most and the second most prevalent nucleotide substitutions in the CNCs (Figure 2A). In tumor samples, while the C:G→A:T transversions were the most prevalent nucleotide substitutions, the C:G→T:A transitions represented the second most prevalent nucleotide substitutions (Figure 2A). In CNC samples, the C:G→T:A transitions were the most prevalent nucleotide substitutions in nonsmokers, current smokers and reformed smokers (Figure 2B). In consistence with previous report^16^, the C:G→T:A transitions were the most prevalent nucleotide substitutions in tumor samples of nonsmokers (Figure 2C), whereas the C:G→A:T transversions were the most prevalent nucleotide substitutions in tumor samples of smokers (Figure 2C).

**Figure 2.**
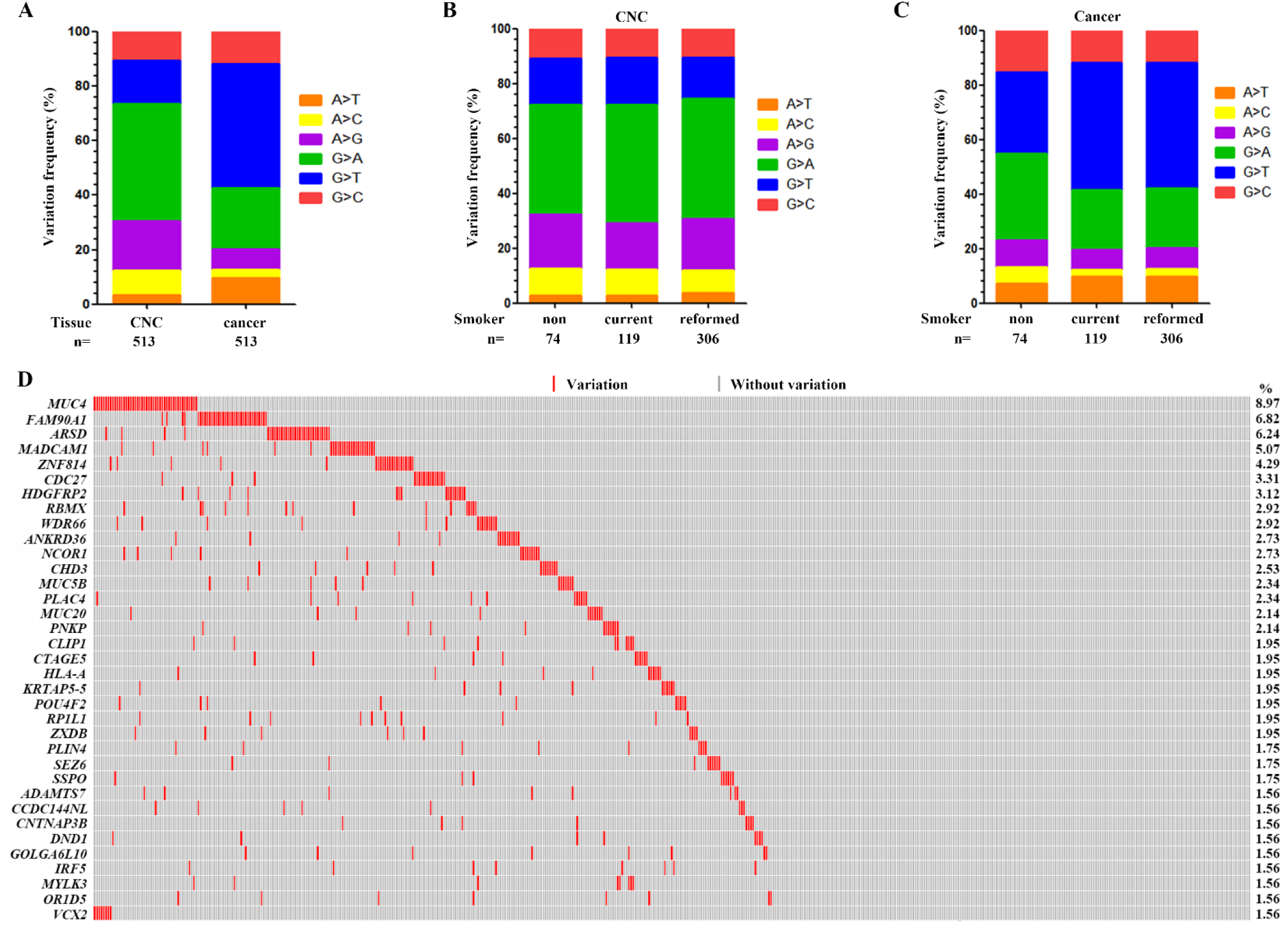
Genomic variations in the CNC samples. (A) The frequency of each type of single base substitution in CNCs and tumor samples of the 513 patients with LUAD. (B) The proportion of nucleotide changes in CNC samples of nonsmokers, current smokers and reformed smokers. (C) The frequency of base substitution in tumor samples of nonsmokers, current smokers and reformed smokers. (D) Significantly altered genes in CNCs of the patients. CNC samples are arranged from left to right in the top track.

### Recurrently altered genes in CNC samples

We reported that there were 16 genes which had a variation rate of more than 2% in the 513 CNCs (Figure 2D and Table S5). *MUC4* represented the most frequently altered gene, which was mutated in 46/513 (8.97%) of the CNC samples (Figure 2D). *FAM90A1*, *ARSD*, *MADCAM1*, *ZNF814*, *CDC27*, and *HDGFRP2* were mutated in 35 (6.82%), 32 (6.24%), 26 (5.07%), 22 (4.29%), 17 (3.31%), and 16 (3.12%) of the 513 CNC samples, respectively (Figure 2D). *TP53*, *KRAS* and *EGFR* which were frequently mutated in LUADs^13^, were mutated in only 2 (0.39%), 0, and 0 CNCs, respectively (Table S5).

*MUC4* encodes an integral membrane glycoprotein found on the cell surface and plays a role in tumor progression^17^. We found 35 types of variations distributed throughout the entire gene of *MUC4*, including 4 recurrent ones (Figure 3). Frequent variations were found in *FAM90A1*^18^, in that in all 35 patients with variant *FAM90A1*, the nucleotide G of codon (ACG) for T344 was replaced by A and a codon for valine (GUC) was inserted in 12:8374781 position at Chr 8, leading to insertion of V right after T344 (Figure 3). There were 67 variations in *Arylsulfatase D* (*ARSD*) gene^19^ in 32 CNCs, with S157F, G175D, and S176K substitutions found in 9 (14.8%), 10 (16.4%), and 10 (16.4%) of the total variations. Twenty-nine alterations were found in *MADCAM1^20^* in 26 CNCs (Figure 3). Variations in *ZNF814*, *CDC27*, *HDGFRP2*, *RBMX*, *WDR66*, and *ANKRD36* in CNCs were also shown in Figure 3.

**Figure 3.**
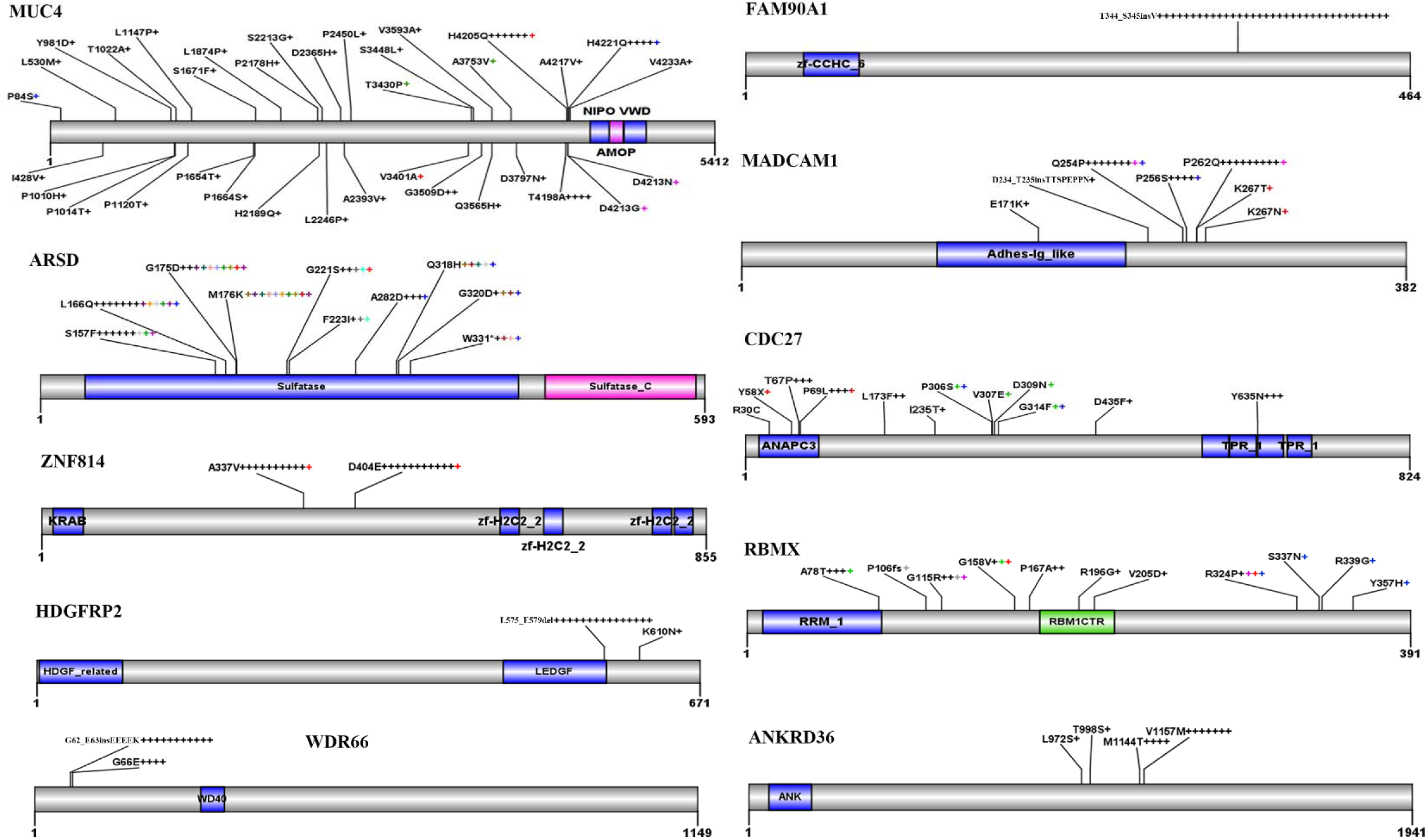
Recurrent variations in some representative genes in CNCs. Schematic representations of proteins encoded by the genes are shown. Numbers refer to amino acid residues. Each “+” corresponds to an independent, altered CNC sample. Variations within a gene indicated by a same-colored “+” are found in the same patient.

### Altered signaling pathways

Pathway analysis was performed using the KEGG (Kyoto Encyclopedia of Genes and Genomes) database, and the results showed that genes involved in type I diabetes mellitus, allograft rejection, graft-versus-host disease, autoimmune thyroid disease, antigen processing and present were the most frequently altered genes (Figure S3A). The GO analysis confirmed that immune response genes were altered in CNC samples, and genes associated heart, muscle, and cell development were also perturbed in CNCs (Figure S3B).

### Variations associated with poor prognosis

We analyzed the potential association between the CNC variations and the prognosis of the patients using the Kaplan-Meier method, and found that variations in 8 genes were associated with the prognosis of 502 patients whose survival information was available (Figure 4). Sixteen variations were found in *MUC5B* in 12 (2.34%) of the 513 patients (Figure 4A). As compared with patients (n=490) harboring wild type *MUC5B*, those with mutant transcript had much shorter overall survival (P=0.013; Figure 4A). *ZXDB*^21^ variations were seen in 10 (1.95%) of the patients, whose overall survival time was shorter than those (n=492) with wild type *ZXDB* (Figure 4A). Patients with variant *PLIN4^22^*, *CCDC144NL*, *CNTNAP3B^23^*, or *CCDC180^24^* in CNCs also had shorter overall survival than patients with wild type transcripts (Figure 4A).

**Figure 4.**
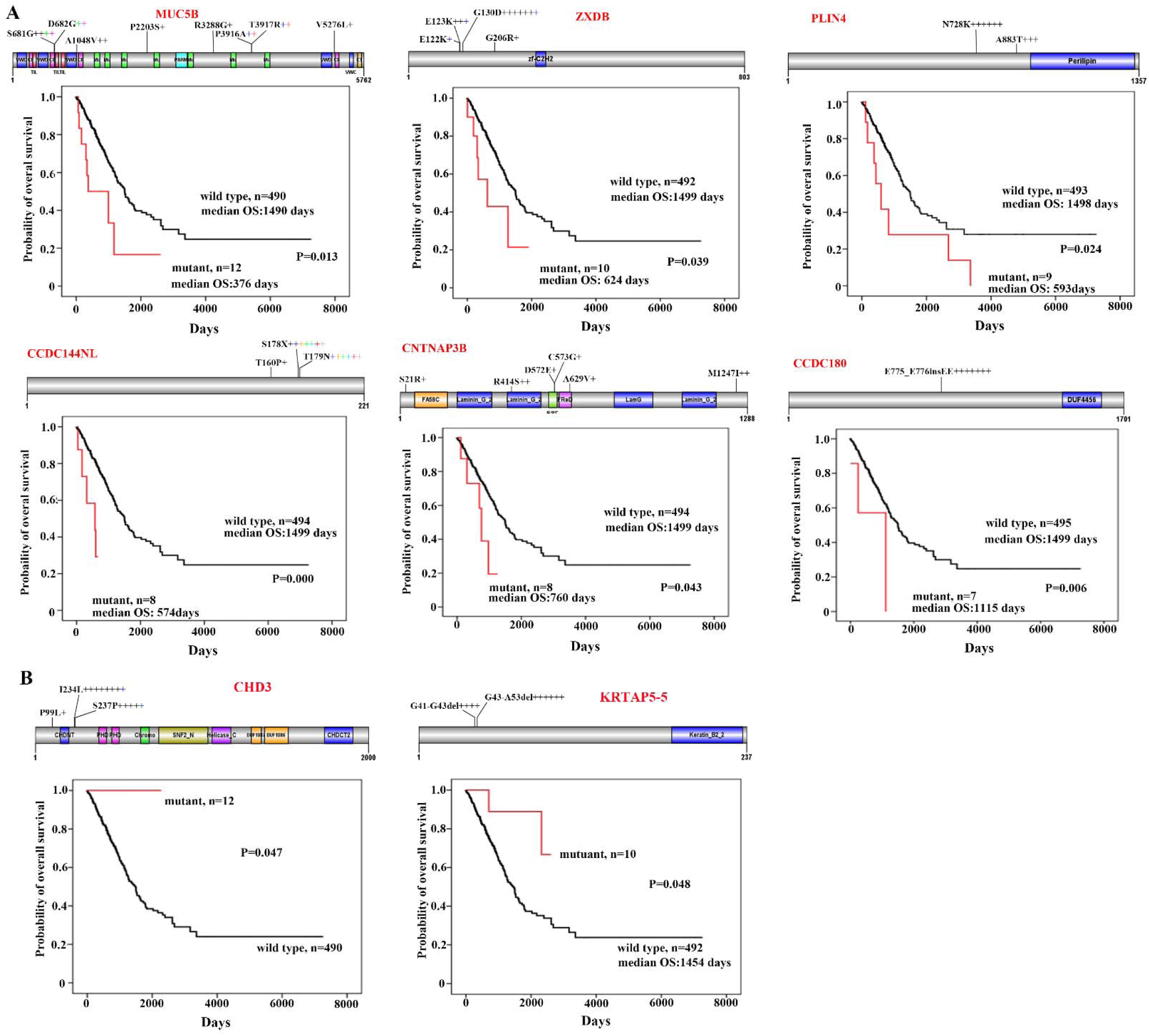
CNC variations associated with prognosis of the patients. (A) Variations of MUC5B, ZXDB, PLIN4, CCDC144NL, CNTNAP3B, and CCDC180 in CNCs and Kaplan–Meier curve for overall survival of the patients with wild type or variant transcripts. Schematic representations of proteins encoded by the genes are shown. (B) Overall survival of patients harboring wild type or variant CHD3 or KRTAP5-5 in their CNCs.

### Variations in CNC samples associated with favorable prognosis of the patients

We found 14 variations in the *Chromodomain Helicase DNA Binding Protein 3* (*CHD3*^25^) gene in CNC samples of 13 (2.53%) of the 513 patients. These variations encoded proteins with P99Lsubstitution in 1 patient, I234L substitution in 8 patients, and S237P change in 5 patients (Figure 4B). Interestingly, none of the variant *CHD3*-bearing patients died within a follow-up of up to 2261 days, whereas the median overall survival of patients with wild type CHD3 (n=490) was 1492 days (range, 0-7248 days) (p=0.047; Figure 4B). Deletion mutations, i.e., deletion of codons for G41 – G43 and G43 – A53 of the protein product encoded by *KRTAP5-5*^26^ gene, were found in CNCs of 10 patients. Of note, patients with variant *KRTAP5-5* had longer overall survival time than those with wild type transcript (p=0.048; Figure 4B).

## Discussion

In our work characterizing somatic mutations in air pollution-related lung cancer^27^, we unexpectedly noted that compared to reference human genome and cancerous tissues, the normal control lung tissues also had single nucleotide variations. This was the impetus for us to conduct this study. We used the “normal-tumor pairs” method to investigate the genomic variations in CNC samples, and reported that the CNCs did harbor genomic alterations that were confirmed by Sanger capillary sequencing of PCR products (Figure 1). These observations were validated in TCGA lung cancer genome data. In CNC genomes of TCGA datasets, the variation rates were 0.2721 exonic variations/Mb and 5.2885 altered genes/sample, and the recurrent variants were of intermediate frequency (~ 8.97% of the patients), whereas variations in 8 genes were associated with poor or favorable prognosis of the patients. We also dissected CNC variations in lung squamous cell carcinoma, and reported a mean of 0.5661 exonic variations/Mb and 7.7887 altered genes/sample in 478 patients. These results uncover the previously unidentified CNC variation, and suggest a role of these alterations in lung carcinogenesis.

Some human carcinogens exhibit specific mutation signatures in the genome. For instance, PAHs which are the main carcinogens of tobacco smoke and air pollution^5, 28^, induce the C:G→A:T substitutions in the genome^7, 15, 16^. The alkylating agent N-methyl-N-nitro-Nnitrosoguanidine (MNNG) which is found in tobacco smoke and a representative molecule of the N-nitroso compounds formed in human stomach^29^, causes the C:G→T:A nucleotide changes in the genome^15^. We found that in CNCs of LUADs of both the smokers and nonsmokers, the C:G→T:A transitions were the predominant nucleotide changes (Figure 2A, B). In tumor samples of the patients, the C:G→T:A substitutions were the predominant nucleotide changes in non-smokers, while C:G→A:T transversions were the most frequently detected substitutions in smokers (Figure 2C). Though C:G→T:A substitutions may be the mutational fingerprints of ageing^30^, these changes in CNCs may be the result of exposure to environmental carcinogens (including second hand smoke in nonsmoker) rather than a consequence of ageing, because in this study the SNPs including those of elder individuals had been filtered out. These data further suggest the key role of environmental factors in lung carcinogenesis.

The significance of these CNC variations in lung carcinogenesis remains unclear. One possibility was that some of the variant genes (e.g., *MUC4*) were pro-oncogenes, cells harboring these CNC variations were in a “precancerous” stage, and accumulation of other mutations would result in transformation and development of malignant neoplasms. Hence, cancer may have multi-clonal origins. Secondly, many of the CNC altered genes were associated with immune response (Figure S3), which may facilitate avoiding immune destruction and cancer initiation. These possibilities warrant further investigation, and CNC variations in other types of cancers including cancers of oral, esophagus, stomach, colorectal, and liver, are worthy of further dissection.

Why CNC mutations were associated with prognosis represents an open question. We hypothesized that these observations may suggest the interactions between CNC tissues and tumor cells, constituting a micro-environment to foster or constrain cancer growth. Those CNC variations associated with poor outcome (*MUC5B*, *ZXDB*, *PLIN4*, *CCDC144NL*, *CNTNAP3B*, and *CCDC180*) may probably facilitate cancer initiation and progression, and may be a new clue to study cancer metastasis and evolution. *CHD3* encodes a chromatin-remodeling factor to repress transcription and is involved in breast cancer^31^. *KRTAP5-5* is an oncogene regulating cancer cell motility and vascular invasion^32^. Whether variations in these two genes modulate transcription machinery or tumor microenvironment to constrain cancer and thus contribute to favorable prognosis, needs to be determined. In addition, how to target the CNC variations to develop novel therapeutics represents another open question.

## Materials and methods

### Patients, whole genome sequencing, and validation by Sanger capillary sequencing

The aim of this study was to address whether there is any genomic alteration in CNCs rather than counterpart tumor tissues (Figure S1). This study was approved by the research ethics committee of our institute. Genomic DNAs were isolated from normal lung tissues (5 cm or more away from the tumors) and adjacent tumors of 11 patients with previously untreated LUAD, the sequencing libraries were constructed, and sequenced using the Illumina Hiseq2000 platform^7^. The genome sequences of counterpart normal lung tissues were analyzed by the Genome Analysis Toolkit (GATK)^33^ UnifiedGenotyper and VarScan, compared with a reference human genome (hg19, downloaded from http://genome.ucsc.edu/)^34^ and filtered against dbSNP138 common germline variants (downloaded from http://genome.ucsc.edu/) and those also detected in tumor samples. Variants from sequencing artifacts were removed by VarScan fpfilter. The following criteria were used to call variations: average base quality ≥ 20, average mapping quality ≥ 20, depth ≥ 6, and variant allele frequency in counterpart sample ≥ 0.20 and ≤ 0.05 in tumor sample. The called variations were validated by polymerase chain reaction (PCR) and Sanger capillary sequencing using genomic DNAs of the patients and primers listed in Table S1. Mutations in cancer genome were also analyzed by the GATK using the above criteria.

### Analyses of TCGA LUAD genome data

The TCGA genome data of 513 LUADs (Table 1) were downloaded from the Cancer Genomics Hub (CGHub) (https://cghub.ucsc.edu/) with approval by the National Institutes of Health (NIH; approval number #24437-4), and analyzed by GATK using the criteria for CNC variants.

### Statistics

All of the values were evaluated using the SPSS 17.0 software for Windows (SPSS Inc., Chicago, Illinois). Statistically significant differences were determined by Student’s *t*-test, and the survival curves were plotted according to the Kaplan-Meier method and compared by the log-rank test. *P* values less than 0.05 were considered statistically significant in all cases.

## Acknowledgements

The results shown here are in part based upon data generated by The Cancer Genome Atlas managed by the NCI and NHGRI. Information about TCGA can be found at http://cancergenome.nih.gov. The dbGaP accession number is phs000178.v9.p8.

## Author Contributions

The project was conceived and designed by GBZ. The experiments were conducted by BZ, LWQ. The genome data were analyzed by LWQ, BZ, GZW, and YC. The manuscript was written by GBZ.

## SUPPLEMENTARY INFORMATION

Supplementary information includes 3 figures and 5 tables.

